# Exploring the Links between Specific Depression Symptoms and Brain Structure: A Network Study

**DOI:** 10.1101/762609

**Authors:** Eva Hilland, Nils Inge Landrø, Brage Kraft, Christian K. Tamnes, Eiko I. Fried, Luigi A. Maglanoc, Rune Jonassen

## Abstract

The network approach to psychopathology has recently received considerable attention, and is as a novel way of conceptualizing mental disorders as causally interacting symptoms. In this study, we modeled a joint network of depression symptoms and depression-related brain structures, using 21 symptoms and five regional brain measures. We used a mixed sample of 268 individuals previously treated for one or more major depressive episodes and never depressed individuals. The network revealed associations between brain structure and unique depressive symptoms, which may clarify relationships regarding symptomatic and biological heterogeneity in depression.

## Introduction

Various patterns of structural brain abnormalities have been associated with depression, yet sensitive, specific and clinically predictive brain correlates have proven to be difficult to characterize[1]. The currently best available empirical evidence on neuroanatomical differences between patients with major depression (MDD) and healthy controls are two meta-analyses of approximately 10.000 individuals[2, 3]. These reports show widespread alterations in cortical regions and in hippocampal volume, but no associations between depression severity and brain structure. Inconsistencies in the neuroimaging literature may be explained by the fact that depression is highly heterogeneous, featuring over 50 symptoms[4], where symptom constellations may reflect different phenomena with distinct underlying biological causes[1].

Understanding the neural substrates of specific symptoms may provide important information about mechanisms underlying depression vulnerability. A growing body of research under the umbrella term ‘network approach’ has recently received considerable attention[5]; the approach understands and aims to model mental disorders as systems of causally interacting symptoms. So far, network studies have been based on symptoms and environmental factors, ignoring relevant neurobiological factors[6]. Here, we address this knowledge gap by modelling a joint network of depression-related brain structures and individual depression symptoms, using 21 symptoms and five regional brain measures. The sample is a mixed group of individuals that previously have been treated for one or more major depressive episodes (MDE) and never depressed individuals, with the goal to model a continuum of depression severity.

## Methods

Depression symptoms were measured using the Beck Depression Inventory (BDI-II). MRI images were obtained from a 3T Philips scanner. Whole-brain volumetric segmentation and cortical surface reconstruction of MRI images was performed with FreeSurfer 5.3 (https://surfer.nmr.mgh.harvard.edu/). Five regional brain measures were selected based on the MDD case-control differences showing the largest bilateral effects in the studies from the ENIGMA MMD working group[2, 3]: hippocampal volume and cortical thickness in four regions - medial orbitofrontal cortex (mOFC), fusiform gyrus, insula and cingulate (weighted average of rostral anterior cingulate, caudal anterior cingulate and posterior cingulate). Brain structure measures were averaged across the left and right hemisphere for each participant, and z-residuals of hippocampal volume (controlling for sex and estimated intracranial volume) were calculated for further analyses. A gaussian graphical model of the 26 variables were computed using the R packages qgraph and bootnet, and the graphical LASSO (least absolute shrinkage and selection operator) was used for regularization. (See Supplementary Information for details on MRI acquisition, MRI processing and network analysis).

## Sample

This sample was drawn from two related clinical trials and a case-control research study conducted at the Department of Psychology, University of Oslo. Informed consent was obtained from all participants before enrolment and their anonymity was preserved. The sample consists of 268 adult participants, 191 with at least one MDE (*M* age = 39.4 [*SD* = 13.2], 132 females, *M* education level (ISCED) level 6.0 [*SD* = 0.9], *M* BDI-II score 14.7 [*SD* = 10.4]) and 77 never depressed individuals (*M* age = 41.9 [*SD* = 12.9], *M* education level 5.7 [*SD* = 1.5], *M* BDI-II score 1.7 [*SD* = 2.9], 50 females). BDI-II sum score range was 0 - 49. A total of 172 subjects had experienced two or more MDE’s. 61 participants were currently using antidepressant medication.

## Results

The symptom-brain network is depicted in *Figure 1A*. All brain structures were positively inter-connected, with regularized partial correlations up to 0.40, see *Figure 1B*. Hippocampus was associated with *changes in appetite sadness, loss of interest* and *irritability*. Insula was associated with *loss of interest in sex* and *sadness*. Cingulate had associations with *sadness, crying* and *worthlessness*. Fusiform gyrus had associations with *crying* and *irritability*. (See stability and centrality indices, S1 and S2)

**Figure 1A:**
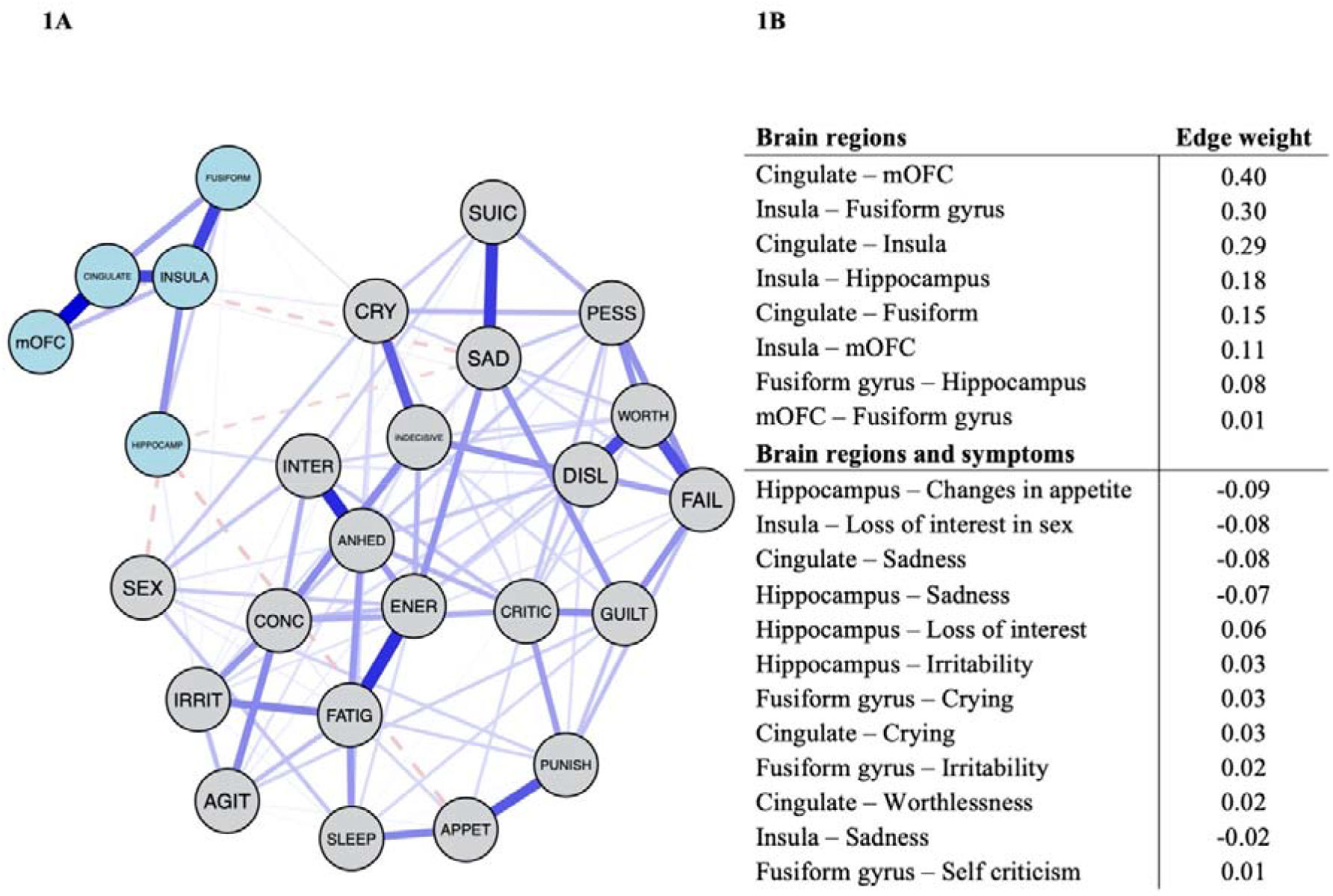
Depression symptom network including five brain areas. Blue lines represent positive associations, red lines negative associations, and the thickness and brightness of an edge indicate the association strength. Label descriptions: mOFC=Medial orbitofrontal cortex, CINGULATE=Rostral-, medial-, and anterior cingulate cortex, INSULA=Insula, FUSIFORM=Fusiform gyrus, HIPPOCAMP=Hippocampus, SAD=Sadness, PESS=Pessimism, FAIL=Past Failure, ANHED=Loss of Pleasure, GUILT=Guilty Feelings, PUNISH=Punishment Feelings, DISL=Self-Dislike, CRITIC=Self-Criticism, SUIC=Suicidal Thoughts or Wishes, CRY=Crying, AGIT=Agitation, INTER=Loss of Interest, INDECISIVE=Indecisiveness, WORTH=Worthlessness, ENER=Loss of Energy, SLEEP=Changes in Sleep Pattern, IRRIT=Irritability, APPET=Changes in Appetite, CONC=Concentration Difficulty, FATIG=Tiredness or Fatigue, SEX= Loss of Interest in Sex. **1B:** Sparse partial correlations between brain structure measures, and between brain structure measures and depressive symptoms in the network model.

## Discussion

Here we establish the first link between individual depression symptoms and neuroanatomy using network analysis. Our results broadly align with prior literature showing that depression symptoms differentially relate to important outcomes such as impairment and risk factors, and demonstrate the importance of studying specific features of depression over one heterogeneous category[5, 6]. The associations between symptoms and brain structure may reflect the heterogeneous nature of the disorder, and may offer important cues about underlying neural mechanisms in MDD. The results await replication in larger samples and other patient groups. In this study depression history was assessed retrospectively and previous MDE was classified independent of type of treatment, combination treatment, treatment response or time since the last episode. We hope the reported results can pave the way for future studies integrating neurobiological measures in network analyses, which represent a step towards validation of biomarkers.

## Acknowledgements

We thank the Department of Psychiatry, Diakonhjemmet Hospital for help and support with recruiting patients, and the Intervention Center, OUS for radiological assistance in MRI protocols, data acquisitions and screening for unexpected neuropathological findings. We thank Tor Endestad for establishing the infrastructure for MRI research at the Department of Psychology, University of Oslo. We want to thank our MRI research assistant Dani Beck. We also thank our external recruitment sites; Unicare, Coperiosenteret AS, Torgny Syrstad, MD, Synergi Helse AS and Lovisenberg Hospital.

